# Simultaneous Electrophysiology and Optogenetic Perturbation of the Same Neurons in Chronically Implanted Animals using μLED Silicon Probes

**DOI:** 10.1101/2023.02.05.527184

**Authors:** Nathaniel R. Kinsky, Mihály Vöröslakos, Jose Roberto Lopez Ruiz, Laurel Watkins de Jong, Nathan Slager, Sam McKenzie, Euisik Yoon, Kamran Diba

## Abstract

Optogenetics are a powerful tool for testing how a neural circuit influences neural activity, cognition, and behavior. Accordingly, the number of studies employing optogenetic perturbation has grown exponentially over the last decade. However, recent studies have highlighted that the impact of optogenetic stimulation/silencing can vary depending on the construct used, the local microcircuit connectivity, extent/power of illumination, and neuron types perturbed. Despite these caveats, the majority of studies employ optogenetics without simultaneously recording neural activity in the circuit that is being perturbed. This dearth of simultaneously recorded neural data is due in part to technical difficulties in combining optogenetics and extracellular electrophysiology. The recent introduction of μLED silicon probes, which feature independently controllable miniature LEDs embedded at several levels of each of multiple shanks of silicon probes, provides a tractable method for temporally and spatially precise interrogation of neural circuits. Here, we provide a protocol addressing how to perform chronic recordings using μLED probes. This protocol provides a schematic for performing causal and reproducible interrogations of neural circuits and addresses all phases of the recording process: introduction of optogenetic construct, implantation of the μLED probe, performing simultaneous optogenetics and electrophysiology *in vivo*, and post-processing of recorded data.

**SUMMARY:** This method allows a researcher to simultaneously perturb neural activity and record electrophysiological signal from the same neurons with high spatial specificity using silicon probes with integrated μLEDs. We outline a procedure detailing all stages of the process for performing reliable μLED experiments in chronically implanted rodents.

## INTRODUCTION

Optogenetics provides a powerful tool for causally testing how a neural activity influences behavior, cognition, and internal neural dynamics. However, optogenetic perturbations can have different effects on neural activity depending on the type of optogenetic construct used, the cellular compartments activated/silenced, and the local circuit anatomy of the perturbed region. For example, silencing cells using the chloride pump halorhodopsin produces rebound spiking after light offset that does not occur with the proton pump Archaerhodopsin (Raimondo et al., 2012). On the other hand, when the same optogenetic constructs are inserted into presynaptic terminals, photoactivation of halorhodopsin reliably reduces neurotransmitter release; yet, for Archaerhodopsin, sustained photoactivation (> 1 min) *increased* evoked transmitter release from baseline due to a pH-change related influx of Calcium (Mahn et al., 2016). These observations illustrate that the biophysical mechanisms through which different constructs work can have opposite effects on neural activity depending on where they are localized. Moreover, local circuit architecture can strongly influence how optogenetic perturbations impacts neighboring cells. For example, activation of single neurons in the hippocampus can suppress or enhance activity in neighboring cells, presumably through interneuron-dependent inhibition (McKenzie et al., 2021; Robinson et al., 2020) (\Watkins de Jong 2023??). Despite these often conflicting and counterintuitive effects, many studies do not record neural activity during optogenetic perturbation, presumably due to the technical difficulty of combining optogenetics with electrophysiology.

While recently developed tools allow all-optical perturbation of neural circuits with combined *in vivo* calcium imaging and optogenetics, such systems are either prohibitively expensive to most labs (Robinson et al., 2020) and/or do not allow for focal optogenetic targeting of individual neurons (nVoke system, www.inscopix.com). Optrodes, which combine electrophysiology with optogenetic fibers, provide one possible solution (Gilmartin et al., 2013; Stark et al., 2014). However, they require the purchase of a light delivery system using lasers or LEDs and must often be customized by hand, a technically challenging procedure. Furthermore, the large footprint of the optical fibers can produce damage beyond that of electrodes which significantly reduces cell yield. µLED probes provide a viable solution for simultaneous optogenetic perturbation and electrophysiological recording from the same neurons with high spatial specificity (Kim et al., 2020; Wu et al., 2015). In a commercially available 32-channel µLED silicon probe configuration, each probe consists of three independently controllable µLEDs on each of the four shanks. This design allows for flexible, focal light delivery while retaining high cells yields due to the integration of the µLEDs with recording electrodes of each shank. Furthermore, the same blue light delivery can allow for neural excitation with constructs such as oChief (Lin et al., 2009) as well as neural silencing with inhibitory rhodopsins such as stGtACR2 (Mahn et al., 2018).

Here, we provide a protocol that addresses all experimental stages for the reliable use of µLED probes. We begin with important considerations for choosing the appropriate optogenetic construct and how to deliver it to the region of interest. We next describe how to prepare the implant and eventual recovery of the µLED probe. Finally, we provide advice for *in vivo* neural excitation/suppression, troubleshooting steps, and best practices for post-processing of neural data. Since drive building and probe implant are covered comprehensively elsewhere (Vöröslakos, Petersen, et al., 2021), this protocol focuses on viral delivery of the desired opsin prior to implant and post-implant *in vivo* optical perturbations with combined electrophysiology.

## PROTOCOL

1. Introduce optogenetic construct to area of interest
  1.1. Viral delivery
    1.1.1. The efficacy of virally induced protein expression can vary widely depending on the amount injected, model species used, virus serotype injected, and brain region targeted (Aschauer et al., 2013; Burger et al., 2004). If you are using a new optogenetic construct, consider doing an expression study to determine how if your viral construct expresses the opsin in your model species/brain region and assess the best dilution to provide robust expression without overexpression and neurotoxicity. See Resendez et al. (2016) for dilution study methodology applicable to any protein of interest.
    1.1.2. Infuse virus of interest
      1.1.2.1. Prepare rodent for surgery using standard techniques based on protocols approved by your institutional animal care and use committee. Perform a craniotomy just large enough to accommodate your infusion needle at the area of interest and stop any blood flow using pressure and cold, sterile saline.
        1.1.2.1.1. IMPORTANT – Matching up your infusion and probe implant coordinates precisely is vital. if you notice that the infusion craniotomy site is no longer visible when you perform your probe implant 2-3 weeks later, we recommend scoring the skull surface with a dental drill to clearly mark your target area.
      1.1.2.2. Perform three injections of 250nL (50nL/min) each at desired site and +/- 250um (total 750nL).
        1.1.2.2.1. Slowly lower needle or pipette to the lowest injection site, leave for 10 minutes to let settle,
        1.1.2.2.2. Inject at 50nL/min and wait 10 mins to allow for diffusion.
        1.1.2.2.3. Raise to next site and repeat steps 1.1.2.2.1-1.1.2.2.2 until finished
      1.1.2.3. If you are performing deep-brain optogenetics, we recommend performing an additional injection of 250nL above your desired site to allow for testing while lowering probe.
      1.1.2.4. Suture animal and provide post-operative care per your approved animal care and use protocol. Typical wait times for full viral expression are 1-2 weeks for mice, 2-4 weeks for rats.
  1.2. Other methods include using transgenic animals expressing optogenetic constructs (English et al., 2017; McKenzie et al., 2021; Valero et al., 2022) or injecting a Cre virus in an upstream region and Flex virus in the region of interest to optogenetically perturb terminals projecting to your recording area. These techniques are discussed in depth elsewhere and will not be detailed here (Mei & Zhang, 2012; Zeng & Madisen, 2012).
2. Build or purchase silicon probe microdrive
  2.1. Build: Plastic 3D printed drive.
    2.1.1. 3D print drive parts for your choice of drive have been made publicly available. See Vöröslakos, Miyawaki, et al. (2021) for further details and https://github.com/YoonGroupUmich/Microdrive for examples.
    2.1.2. Build drive according to Vöröslakos, Miyawaki, et al. (2021) or instructions provided at https://github.com/YoonGroupUmich/Microdrive.
    2.1.3. Note that drive designs are continuously refined. Inspect the open-source repositories listed above for the most up-to-date designs.
  2.2. Purchase or Build: Metal drive
    2.2.1. Highly stable metal drives with a smaller footprint and similar weight to the plastic drives listed above are also available for building or purchase from 3DNeuro (Vöröslakos, Petersen, et al., 2021)
3. Attach probe to drive
  3.1. See (Vöröslakos, Petersen, et al., 2021) Fig 1 – Video 2 for a video demonstrating the process of attaching the probe to the drive.
4. Test probe prior to implant
  4.1. Verify impedances using Intan recording software or other devices.
    4.1.1. Plug in 32-ch amplifier and connect to recording cable. NOTE: Hold probe electronic interface board (EIB) carefully while plugging/un-plugging headstage and stimulation cable to avoid breaking probe shanks!
    4.1.2. Test impedances using Intan recording software or other devices. Note that impedances will drop following implant into brain tissue.
      4.1.2.1. Submerge mounted probe into saline, connecting a wire that is glued to the side of the container to the ground and reference wires of the probe.
  4.2. Verify µLED functionality (see below on options for powering µLEDs) using OSC1lite board and software.
    4.2.1. Carefully plug in stimulation cable to the OSC1lite **(Figure 1)** and probe EIB (**Figure 2)**. See below for more detailed instructions on using the OSC1lite. We recommend attaching the probe to a stereotax arm to avoid breakage during testing.
    4.2.2. Consult the manufacturer-provided I-V curves and provide current at just below and just above the onset current to verify LED activation. Do not exceed 100μA.
      4.2.2.1. NOTE: While you can use a voltage-controlled waveform generator to power the µLEDs, we do not recommend doing so due to the non-linearity of the I-V curve. If you do use a voltage generator, be especially careful to keep your voltage at levels well below the max shown on the I-V curve and/or use a series resistor to limit your current output **(Figure 3)**.
    4.2.3. Verify activation of each μLED by eye, taking notes of any differences in activation level between μLEDs **(Figure 7)**.
  4.3. The above steps are especially important to perform for new users as it forces the user to become comfortable with plugging/unplugging everything, coordinating two cables, etc. It also allows for identifying and troubleshooting any external sources of electrical noise PRIOR to implanting in a live animal.
5. Implant μLED probe
  5.1. The probe implant process is detailed in full elsewhere: refer to Vandecasteele et al. (2012) for a comprehensive protocol covering probe implant for chronic, in vivo recording in freely moving rodents.
    5.1.1. We highly recommend providing a separate ground and reference screw to help mitigate any line noise issues that might occur (see Discussion).
  5.2. Here we provide µLED specific recommendations for each user to consider based on their region of interest, since drive design and implant procedure are strongly influenced by the targeted brain region.
    5.2.1. For initial implants or when performing multiple implants in the same animal, we recommend using a plastic crown base with copper mesh for implant protection as described in Vöröslakos et al. (2021), to provide maximum flexibility during implant.
    5.2.2. Once the process for a single implant is streamlined, we recommend using a hybrid plastic/mesh cap for mice and fully plastic cap for rats as described in (Vöröslakos et al., 2021)
    5.2.3. NOTE: the application of UV light to cure epoxies can penetrate the skull and photoactivate blue-light opsins such as channelrhodopsin or stGtACR2.
6. Lower probe to region of interest.
  6.1. Gradually move the probe drive down to the target region, keeping track of microdrive screw turns to estimate probe depth (280 µm per turn for drives listed above in section 2) and verifying probe location with electrophysiological signatures specific to the brain region where the probe is implanted.
    6.1.1. Turn 150-300 µm each day maximum at first, then reduce to 75-150 µm as you approach the region of interest.
    6.1.2. NOTE: you should observe an immediate change in neural signal when turning, followed by a slower change over the next several hours as the brain tissue settles around the probe. We recommend waiting at least 6-8 hours before turning the screw again to ensure that you don’t drive the probe past the target region.
  6.2. If you are performing deep brain recordings, we recommend testing the µLEDs in superficial brain regions as you advance the electrodes in order to determine initial illumination settings and identify/troubleshoot any issues prior to reaching your target region. See section 7 for full details.
7. Stimulate/Silence neural spiking activity with OSC1lite
  7.1. Follow all directions for operating OSC1lite at: https://github.com/YoonGroupUmich/osc1lite, most importantly:
  7.2. Tape stimulation and recording cables together to keep clean and facilitate easy connection/disconnection in case of excessive twisting during experimentation.
  7.3. Connect TTL output pins on OSC1lite to event/TTL ports on your recording system. See **Figure 8**.
  7.4. If performing closed-loop or automated, open-loop stimulation/silencing, connect TTL input pins on OSC1lite to hardware supplying the triggering signal.
  7.5. Initialize OSC1lite to probe connection.
    7.5.1. Power on the OSC1lite.
    7.5.2. Connect to open-source OSC1lite software and make sure all channels are off. Make sure you select the device whose last three characters match the identifying characters on your OSC1lite board (**Figure 4 and 5)**.
    7.5.3. Connect 18-pin Omnetics cable to board, then to probe **(Figure 2)**.
    7.5.4. Connect OSC1lite ground to recording system ground.
    7.5.5. Attach 32 channel headstage to probe.
    7.5.6. Connect recording cable (12-pin SPI for Intan) to recording system and then to headstage.
      7.5.6.1. Note that the headstage will block access to all but one side of the stimulation cable connector on the probe. Therefore, we recommend connecting the stimulation cable first. One strategy to deal with excessive cable twisting (e.g., due to longer recordings or working with rats instead of mice) is to build a short 18-pin cable stub approximately the length of the headstage to facilitate easy cable connection/disconnection mid-recording.
    7.5.7. Titrate light levels for stimulation/silencing.
      7.5.7.1. We recommend performing these steps in the animal’s home cage or a dedicated rest box prior to performing any experiments.
      7.5.7.2. Create your desired stimulation waveform in OSC1lite and set appropriate minimum light levels for each µLED based on probe testing protocol outlined in step 4. **See Figure 6**.
        7.5.7.2.1. We recommend a square waveform with 2ms rise time, 100ms pulse width, 2 second period, and 10 pulses to start (**Figure 9)**. This will give you sufficient trials to evaluate the efficacy of stimulation/silencing at a given light level.
      7.5.7.3. Trigger each µLED manually while recording.
        7.5.7.3.1. Adjust light level up/down as your experiment requires to provide the lowest light power necessary for the required level of optogenetic stimulation/suppression. See **Figure 10** for example of a successful stimulation of neurons located near shank 1.
        7.5.7.3.2. Make note of the light level for each µLED and save these as separate waveforms in the OSC1lite software.
      7.5.7.4. Create waveforms for each µLED based on titration results.
        7.5.7.4.1. Make sure to choose the appropriate waveform, trigger source (PC or External), and Mode (One-shot of Continuous) for each µLED in the rightmost window of the OSC1lite software. See **Figure 6 and Figure 9**.
        7.5.7.4.2. Check “Trigger Out” for each µLED whose activation you want to record.
        7.5.7.4.3. Note that L1 occurs at the *top* of the software window for each shank while it is located at the *bottom* of each shank.
  7.5.8. Stimulate/Silence
    7.5.8.1. Option 1: Trigger manually
      7.5.8.1.1. Click “PC Trigger” for individual µLEDs of “Trigger All” to illuminate all µLEDs simultaneously
    7.5.8.2. Option 2: Trigger externally (for automated or closed-loop designs)
      7.5.8.2.1. Make sure to select “External Trigger” in the right window.
      7.5.8.2.2. Send a TTL to the OSC1lite TTLin port(s) from external hardware. 7.5.8.2.2.1. Note that the OSC1lite will apply the appropriate waveform at the *onset* of each external TTL and that the full waveform will occur and will not necessarily match the length of the external TTL.
  7.5.9. Patterned stimulation (Vöröslakos et al., 2022)
    7.5.9.1. For more complex signals, one can use an Arduino to trigger specific µLEDs using the TTL inputs of OSC1Lite.
    7.5.9.2. Set Trigger to external mode, specify current amplitude, duration and duty cycle.
      7.5.9.2.1. NOTE: This preconfigured stimulation waveform will be triggered every time OSC1Lite receives a TTL input.
    7.5.9.3. To test pattern, use Plexon Headstage Tester Unit.
  7.6. Alternative µLED powering options
    7.6.1. µLEDs can also be powered with an on-head current driver design is available (Tarnavsky Eitan et al., 2021)
    7.6.2. Plexon’s NeuroLight Stimulator System is a commercial μLED driver available for purchase.
    7.6.3. Voltage driver. While constructing a simple voltage driver or using a waveform generator to activate µLEDs is possible, we discourage its use and include a discussion here only as a word of caution: unlike with the OSC1 lite or current driver listed above, current and power rise in a sharp, non-linear fashion with small increases in voltage that can easily overpower and burn out a µLED, see **Figure 3**.
      7.6.3.1. To avoid damaging µLEDs, do not exceed the minimum voltage necessary to activate each µLED that you determined during testing (see Step 4).
      7.6.3.2. To minimize stimulation artifact, maintain a voltage at just under (0.1-0.2V) below the minimum level needed to activate each µLED and then step up the voltage to the minimum level to activate the µLED.
8. Probe recovery and post-recovery care
  8.1. Probe recovery and cleaning steps are detailed fully in Vöröslakos et al. (2021). Note that using an enzymatic cleaner such as Ultrazyme is highly recommended as other cleaning agents can compromise µLED and electrode integrity.

## REPRESENTATIVE RESULTS

Following the above protocol will ensure a high probability of successful neural perturbation with µLED probes. Successful focal silencing using all µLEDs on one of the edge-most shanks should result in almost complete neural suppression on that shank, partial inhibition of spiking on the adjacent shank, and no disruption of activity on the furthest shank away **(Figure 11)**. Likewise, successful photoactivation will induce spiking on one shank only and have little to no effect of spiking on adjacent shanks (**Figure 10**). Nonetheless, on occasion no silencing or photoactivation may occur during light stimulation. The two most frequent causes of failed optogenetic perturbation are 1) improperly functioning µLEDs, and 2) a lack of opsin expression near the recording site. Carefully following all steps listed above, including injecting opsin above the region of interest for testing and troubleshooting prior to recording, and carefully testing all µLEDs prior to implant, will mitigate issue #1. Multiple viral injections and/or using transgenic animals can help mitigate issue #2 and performing careful histology following failed stimulation/silencing can aid in troubleshooting any targeting errors as well.

**Figure 1:**
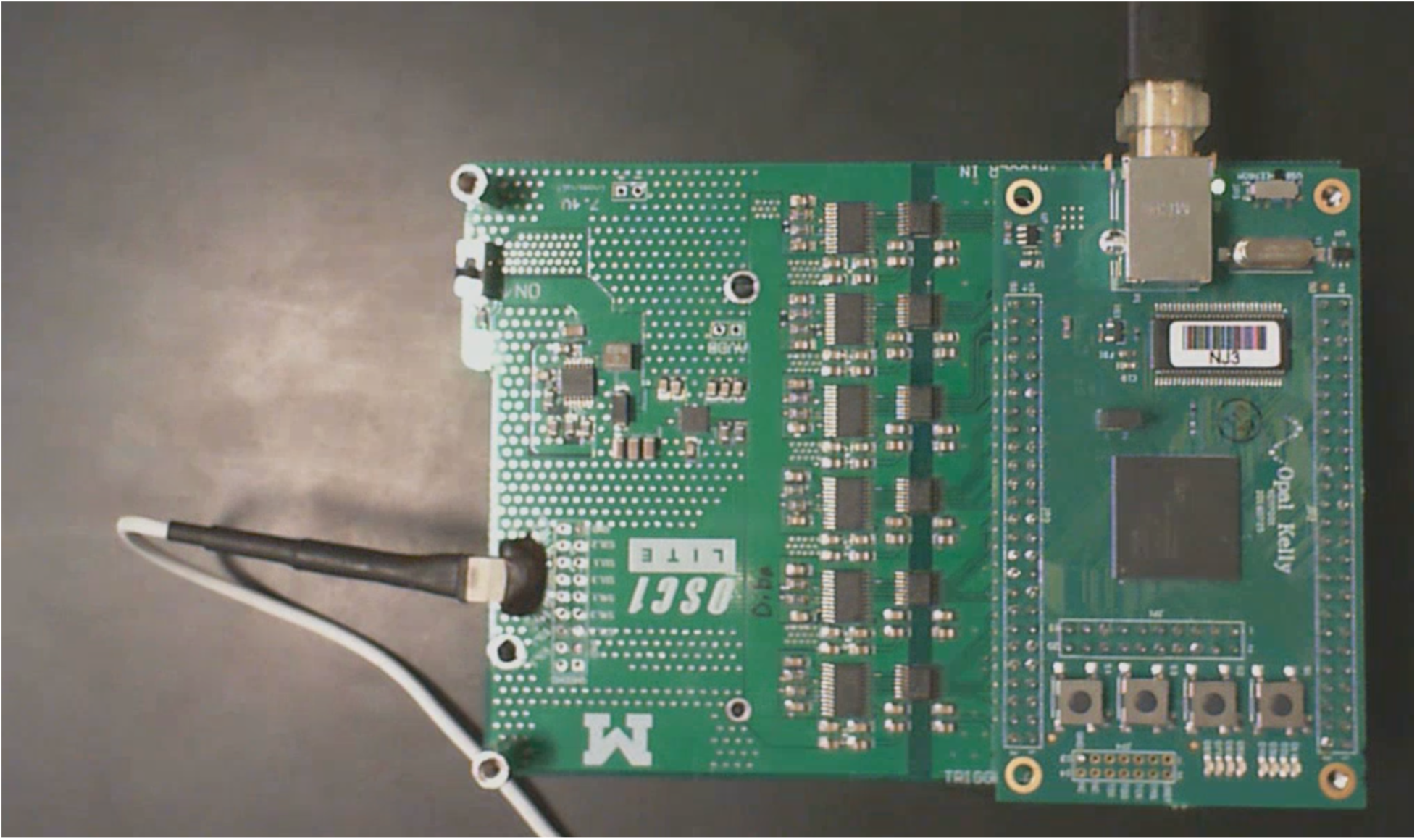
Cable Connections to OSC1lite. 16 pin white stimulation cable shown plugged in on left side of board. USB cable to computer shown at top. Note the “NJ3” identifier just below the USB connection port.

**Figure 2:**
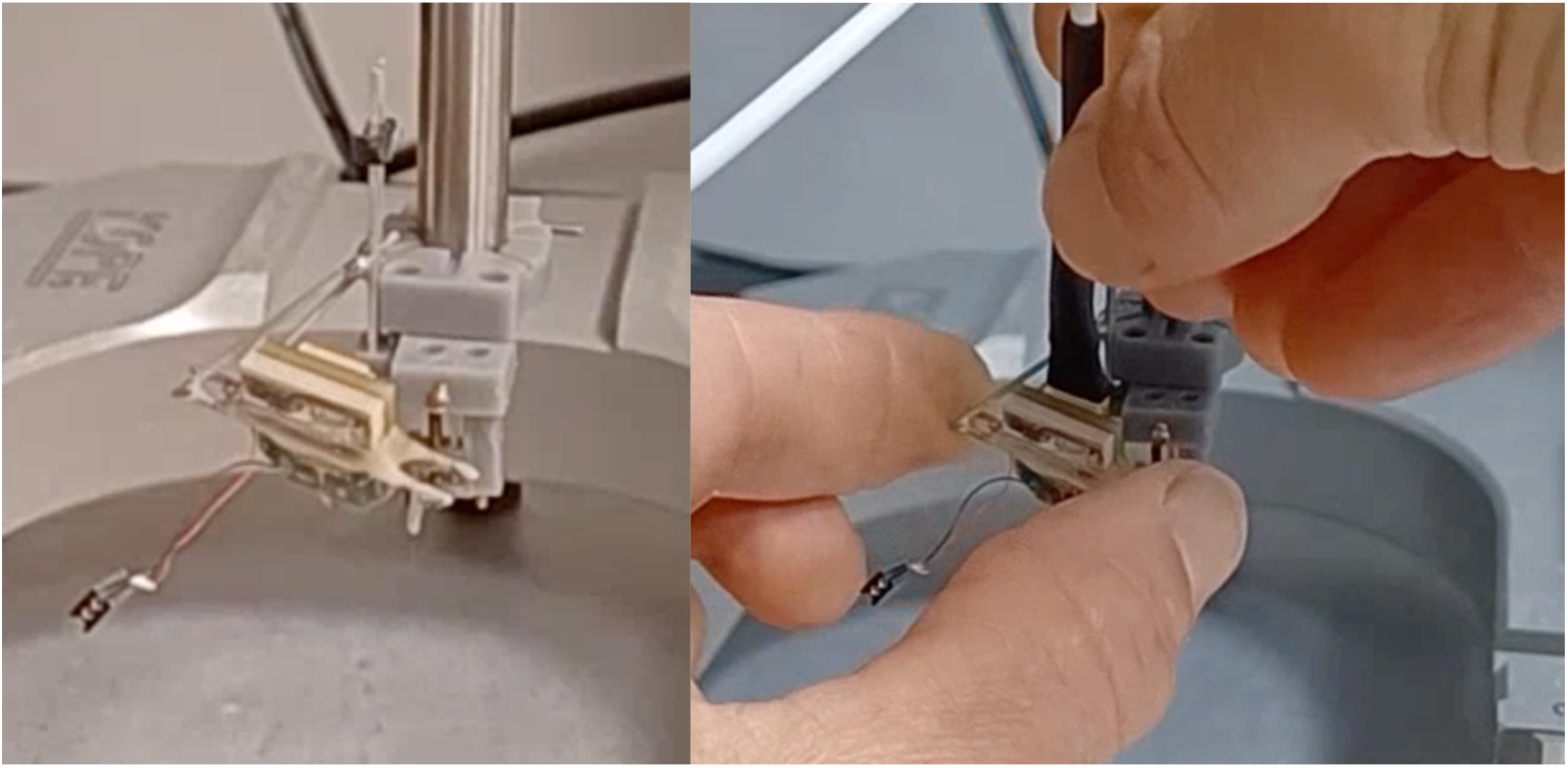
Probe testing configuration. *Left*: μLED probe attached to plastic drive, mounted on stereotax arm for testing prior to implant. *Right:* proper procedure for connecting stimulation cable, note fingers carefully supporting μLED electric interface board during cable attachment.

**Figure 3:**
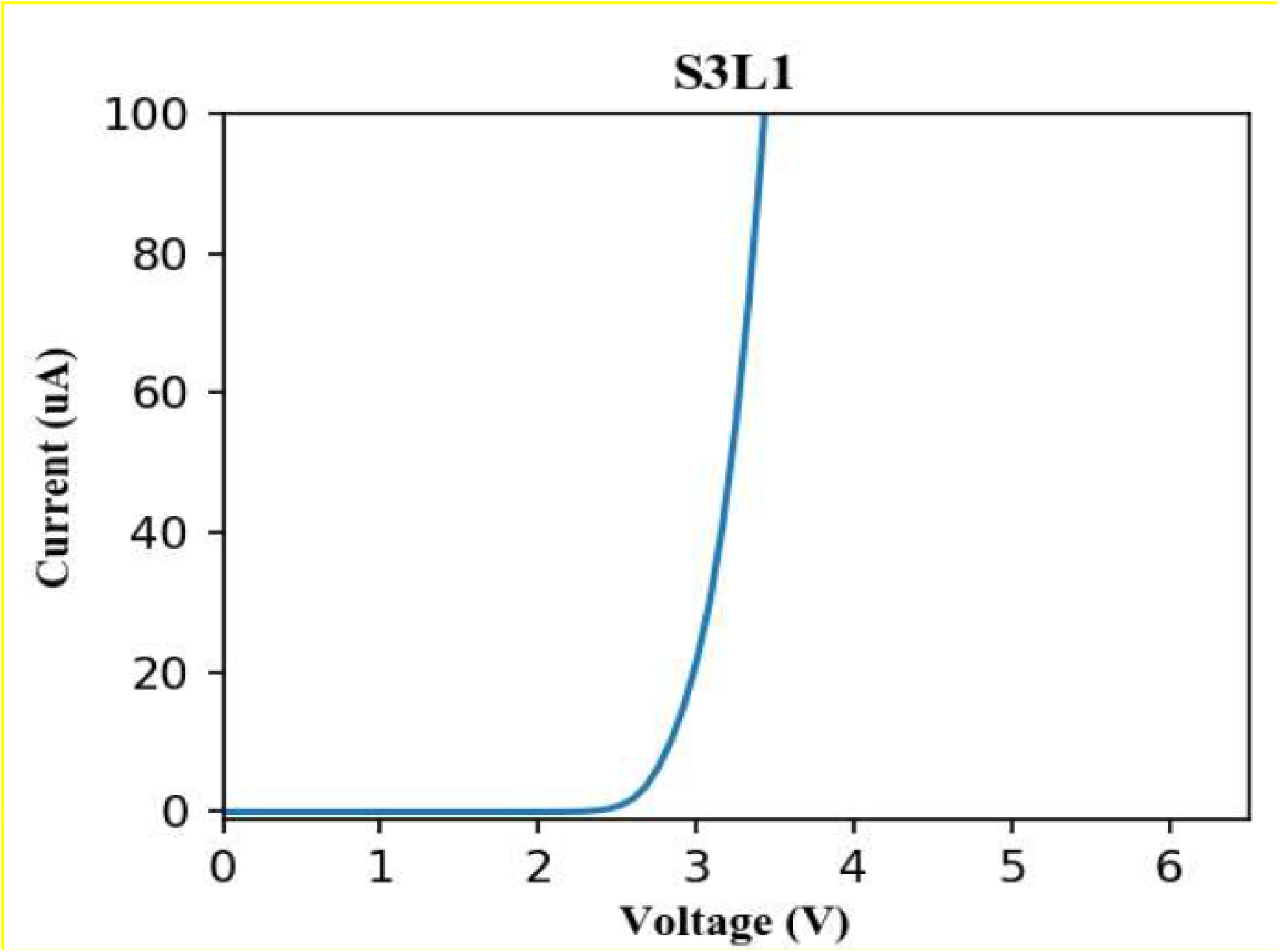
Example I-V curve for μLED 1 (bottom) of shank 3. Notice a sharp, non-linear increase in current at around 3V, highlighting the danger of using voltage drivers to activate μLEDs.

**Figure 4:**
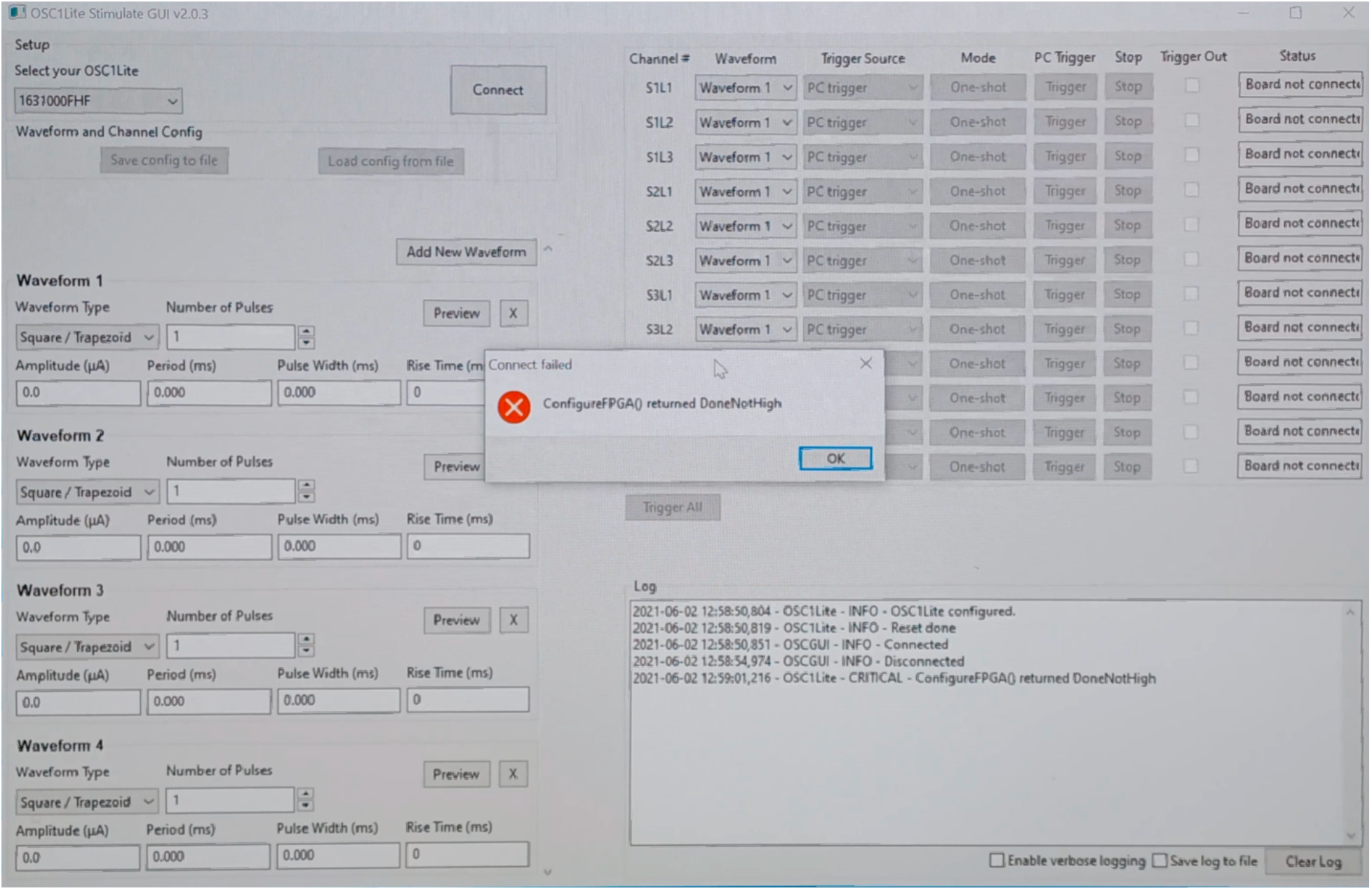
Incorrect selection of OSC1lite in software. Note that the last three characters (“FHF”) do not match the OSC1lite identifier “NJ3” (see Figure 1), resulting in a connection error.

**Figure 5:**
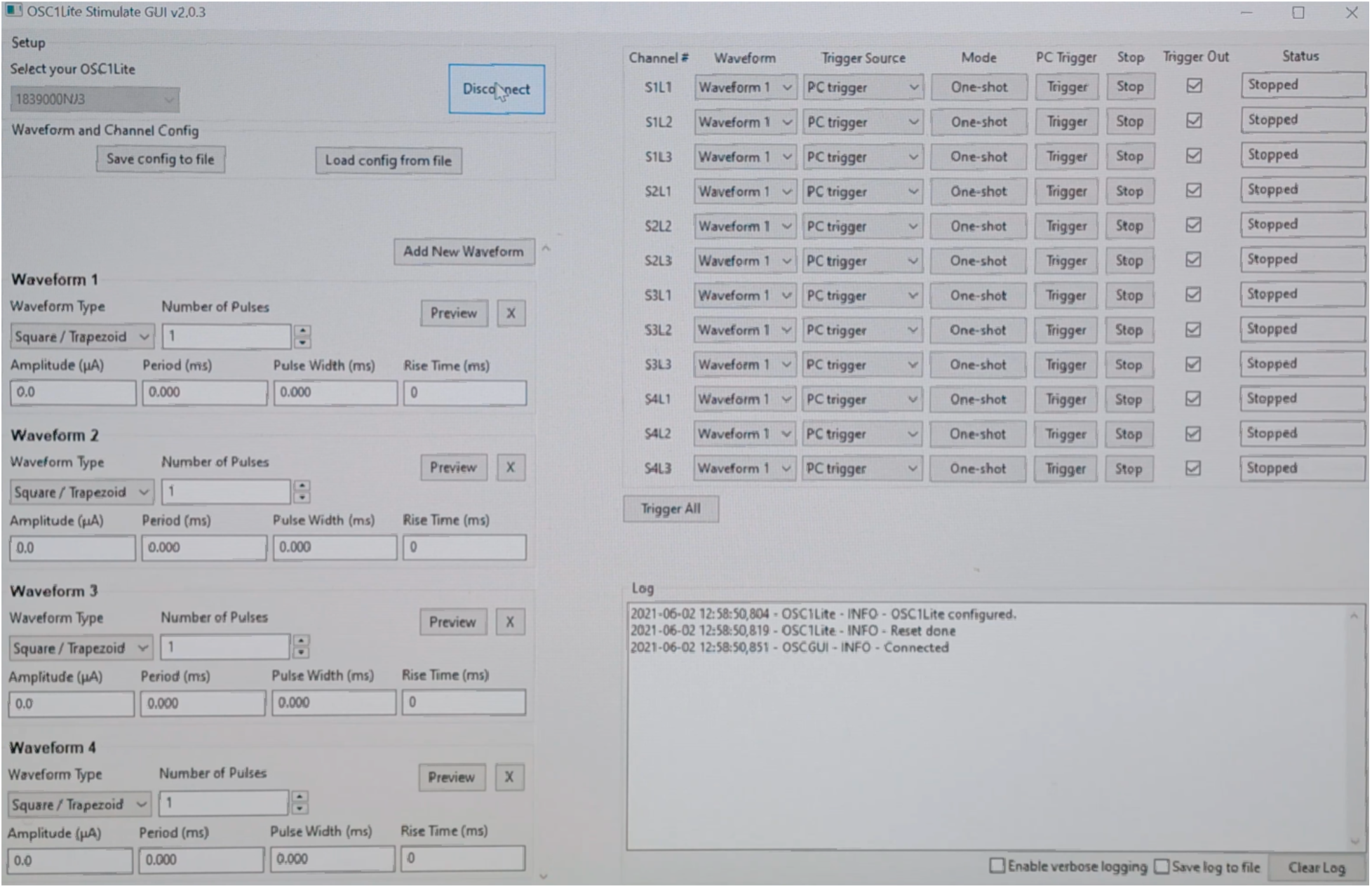
Correct connection to OSC1lite in software. Note that last three digits under “Select your OSC1lite” match the “NJ3” identifier from Figure 1 resulting in a “Connected” printout to the Log in the lower right.

**Figure 6:**
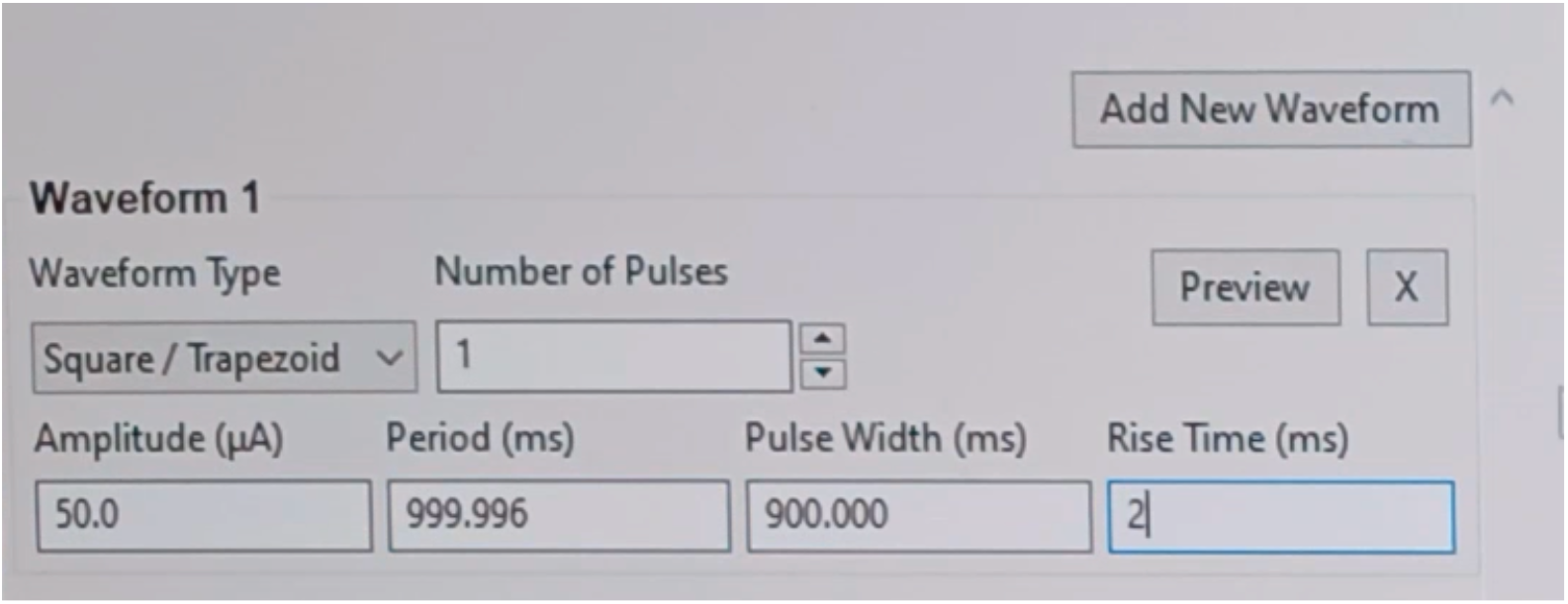
Waveform creation. This figure shows how to create an example waveform which will deliver a single 50 μA pulse lasting 900ms with a 2ms rise time and 2ms fall time to mitigate stimulation artifact. Clicking “Preview” will plot out the waveform created as a visual check.

**Figure 7:**
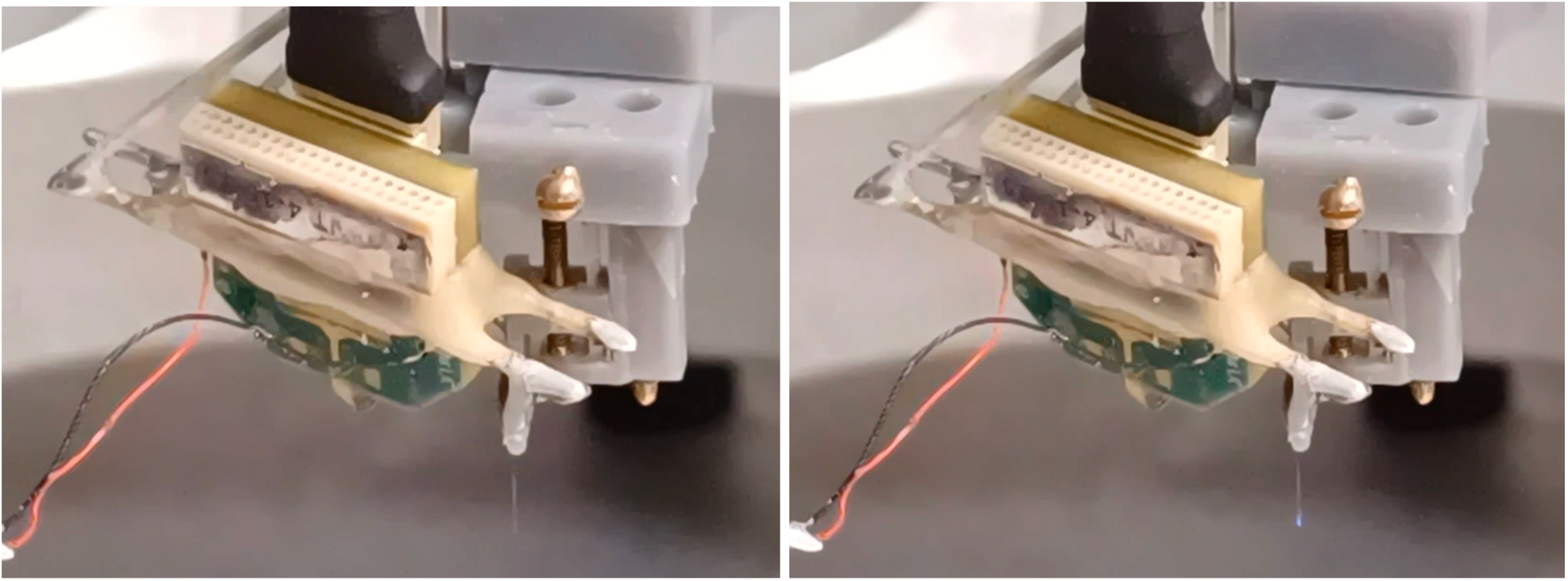
Example of proper LED activation by eye during testing. *Left:* μLED is not active – current level is either too low to activate the light or μLED/OSC1lite software is not functioning properly. *Right:* Proper activation of a μLED by OSC1lite software proper is indicated by a small amount of blue light visible by the naked eye.

**Figure 8:**
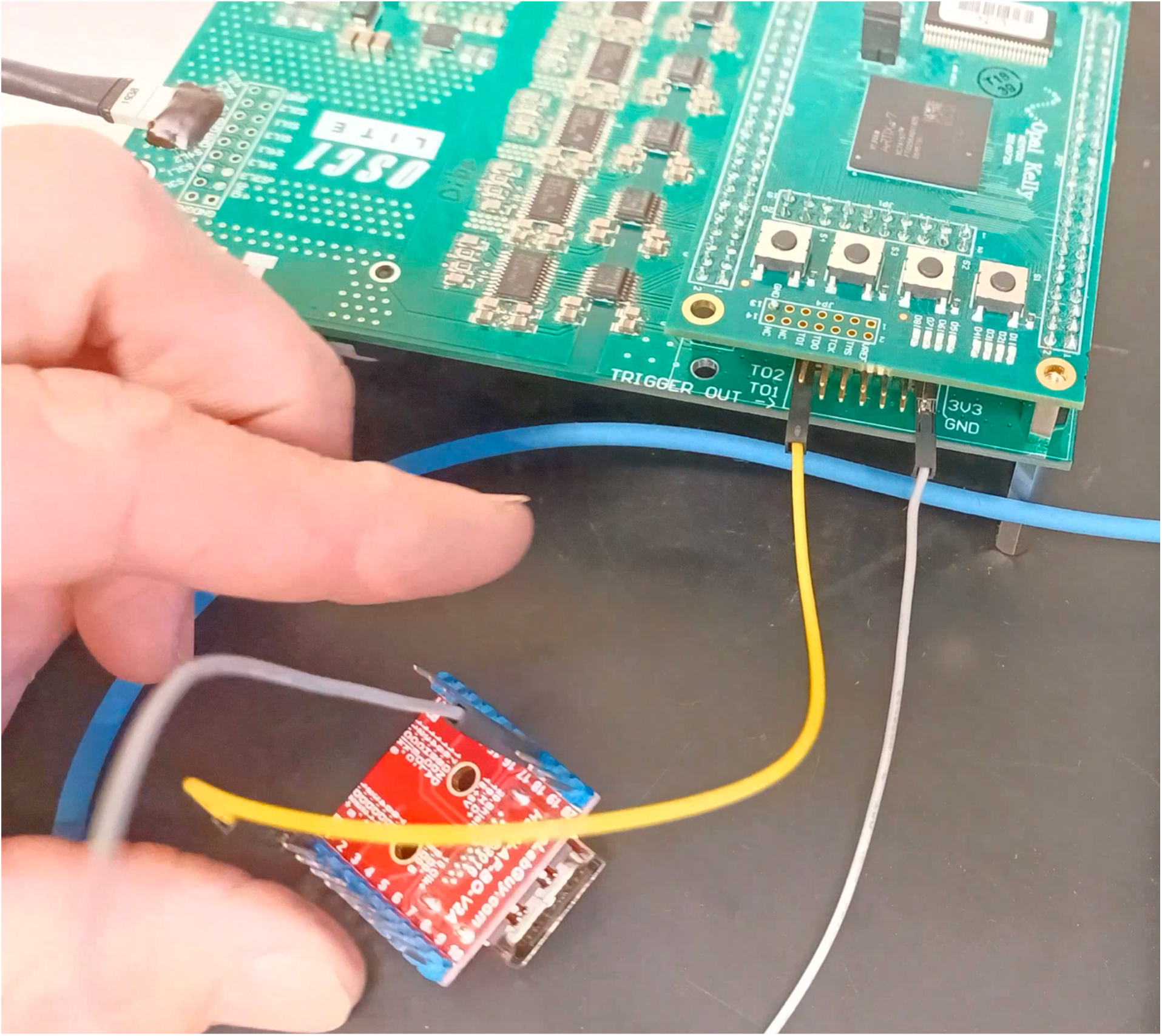
Example OSC1lite Trigger Out port connection. In this case, the shank 1 L2 μLED trigger out port is connected to the D2+ pin and the ground pin port is connected to the ground pin of an HDMI connector board which will be used to transfer a TTL for each activation of the Shank 1 L2 μLED to digital channel 1 of an OpenEphys acquisition system. Make sure the “Trigger Out” box (enabled by default) is checked for each pin you connect to in the OSC1lite software. Similar connections can be made for each μLED, and a similar strategy can be used to collect external triggers to activate μLEDs. Be sure to select “External Trigger” under the “Trigger Source” column in the right side of the OSC1lite software for external triggering.

**Figure 9:**
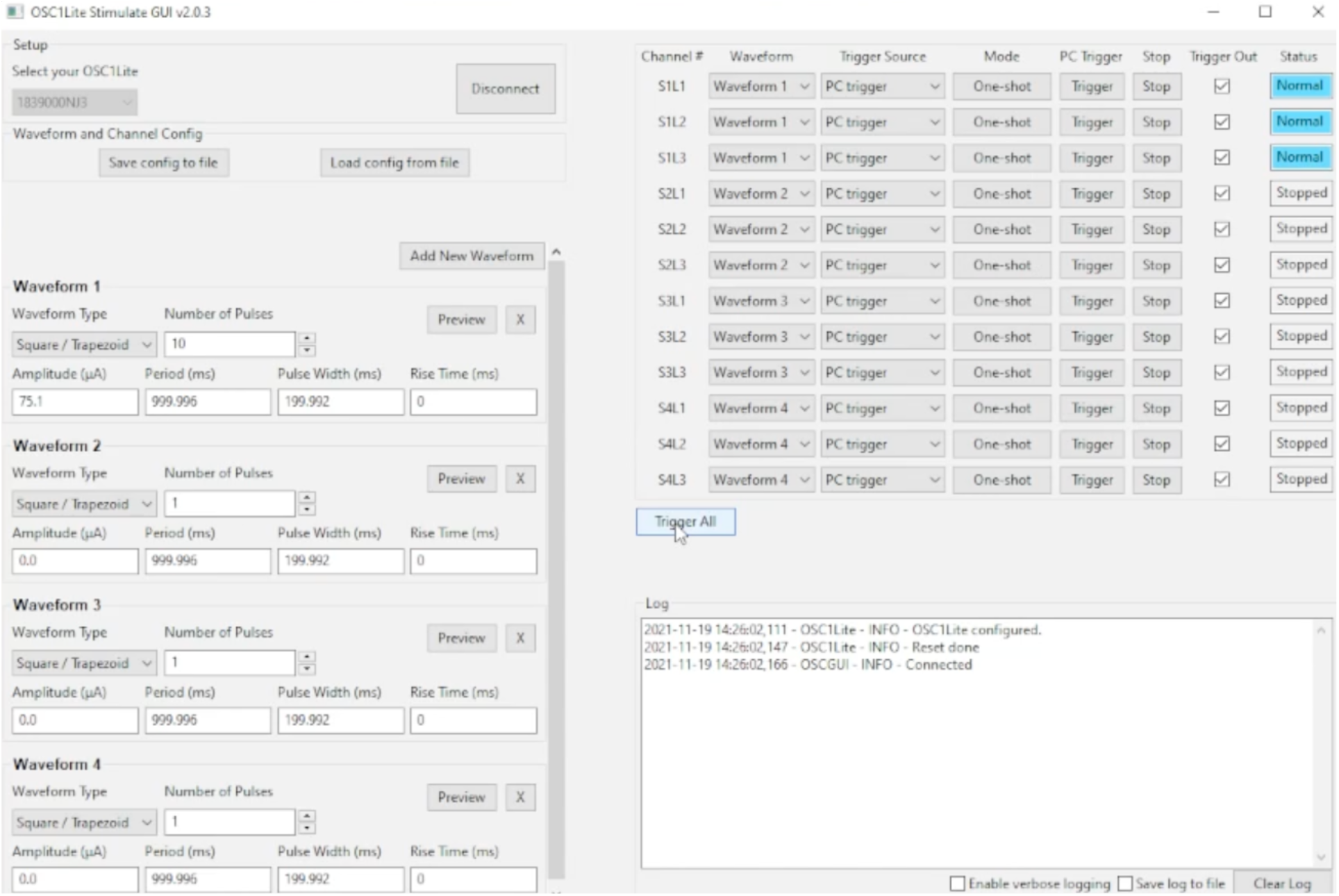
Example in vivo stimulation setup for focal stimulation of neurons near Shank 1 only. Waveform 1 is set to deliver a 200ms, 75µA pulse once every second, 10 pulses total. Waveform 1 is applied to Shank 1 only on the right side of the software GUI such that only Shank 1 LEDs show up in blue as “Normal” during operation while the other LEDs, which are not activated (Amplitude = 0µA), show up as “Stopped”.

**Figure 10:**
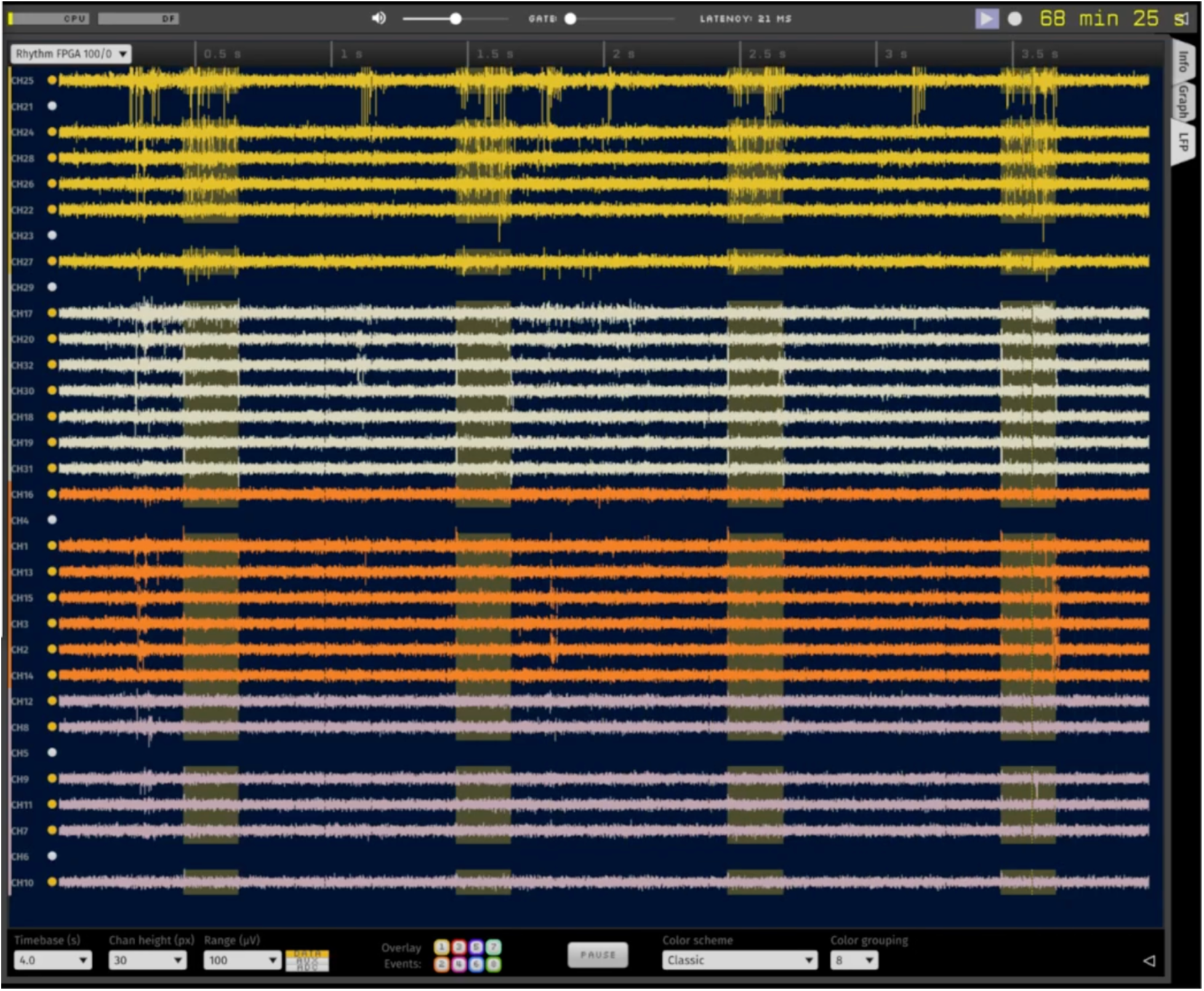
In vivo evoked spiking on Shank 1 using channelrhodopsin. High pass filtered signal recorded with a chronically implanted µLED probe in a rat expressing CHR2 in region CA1 of the hippocampus (high impedance channels are not shown, n = 6 channels). Waveform 1 activation (see Figure 9) evokes robust spiking activity on Shank 1 (yellow) but not on other shanks. Activation times are clearly shown in OpenEphys acquisition software as yellow vertical boxes, logged via sending the Shank 1 Trigger Out pins to Digital Input 1 (see Figure 8).

**Figure 11:**
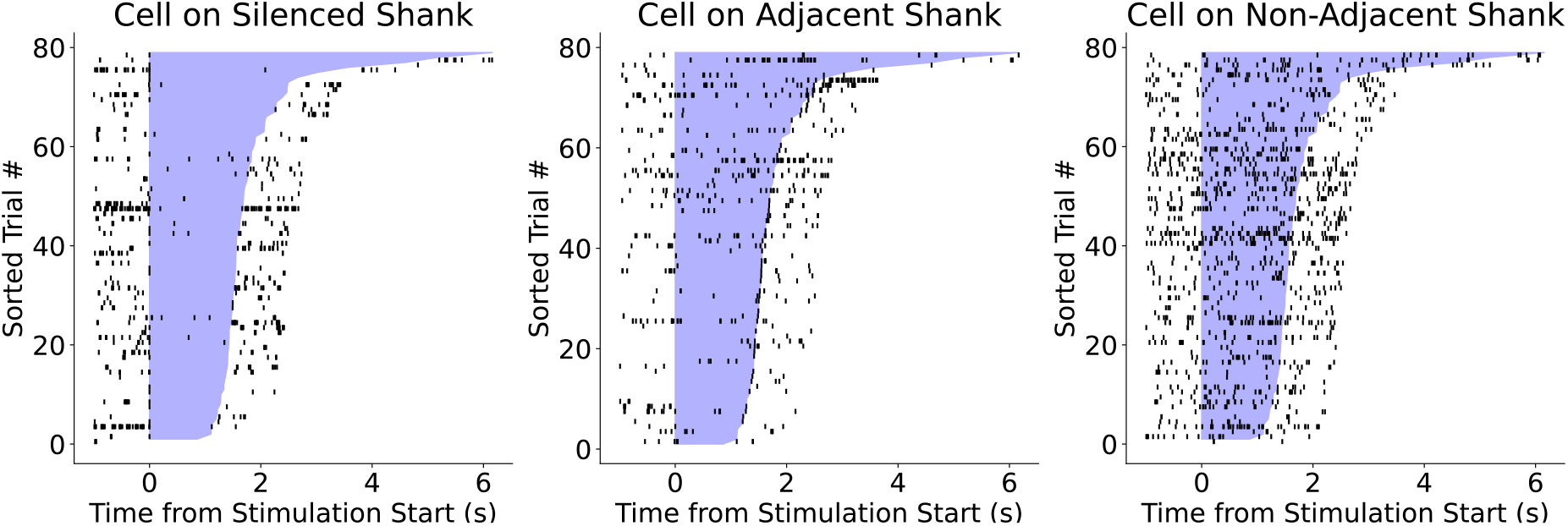
Example of successful, localized silencing of neural activity. Raster plots of single units in the hippocampus during light application on Shank 1 µLEDs (left) only using the inhibitory opsin StGtACR2. Mild suppression is still evident on the adjacent Shank 2 (middle) while spiking activity remains un-perturbed on Shank 3 (right).

## TABLE OF MATERIALS

| Name of Material/<br>Equipment | Company | Catalog<br>Number | Comments/Description |
| --- | --- | --- | --- |
| AAV1-hSyn1-SIO-stGtACR2 | Addgene | 105677 | For optogenetic silencing |
| ChR2 construct | Addgene | 26969 | For optogenetic excitation |
| μLED probe | Neurolight<br>or MINT<br>program | N1-F21-O36/18 | 32 channel silicon probe (4 shanks) with 3 independently controllable μLEDs on each shank.<br>https://github.com/YoonGroupUmich/osc1lite |
| OSC1lite | MINT<br>program | N/A | For controlling μLEDs:<br>https://github.com/YoonGroupUmich/osc1lite |
| 1028 ch Recording system | Intan | RHS2000 | μLEDs are also compatible with many other recording systems, including Plexon models, the OpenEphys acquisition board, and other Intan controllers. See below. |
| <u>Other stimulation and recording options not used in the protocol</u> |  |  |  |
| OmniPlex Neural Data Acquisition System | Plexon | N/A | For recording from μLED probe. |
| OpenEphys Acquisition Board | OpenEphys | N/A | For recording from μLED probe. |
| Neurolight Stimulator System | Plexon | N/A | For controlling μLEDs. |

## DISCUSSION

Here we provide a comprehensive plan for simultaneous optogenetic perturbation of and recording from single neurons. Each of these steps can be important for obtaining optimal performance of μLED probes. First, initial testing prior to *in vivo* activation is vital to ensuring all µLEDs are functioning properly and all electrode impedances are appropriate. Testing is likewise crucial before re-using a μLED probe to verify no damage was done during probe recovery – do not stimulate using any inoperable µLEDs as they can cause extreme artifacts in your recordings. Testing prior to implant also affords the user the ability to practice activating µLEDs, connecting the probe, and interfacing with the software in a controlled environment. The second critical step is the implant surgery. In the case of viral delivery of the opsin of interest, special care must be taken to target the same region where the opsin is expressed. Performing multiple viral infusions offset from one another greatly aids in ensuring the electrodes and opsin end up in close proximity. Care must also be taken to ensure a solid ground and reference screw connection. While many users obtain quality signal using a single screw, having a separate reference wire can help mitigate electrical noise in recordings.

Last, care must be taken to choose the correct opsin for the experimental protocol. Repetitive stimulation with channelrhodopsin can result in reduced efficacy; oChief is a better construct for long periods of continuous stimulation (Hass & Glickfeld, 2016). Similarly, while a number of inhibitory opsins are effective at suppressing neural spiking activity, long bouts of photoinhibition using chloride pump activating constructs, like halorhodopsin, cause rebound excitation upon light offset (Raimondo et al., 2012). Likewise, extended illumination of constructs which utilize proton pumps, like Archaerhodopsin-3, induces paradoxical increases in neurotransmitter release (Mahn et al., 2016). For paradigms requiring extended periods of neural inhibition, a construct such as StGtACr2, which avoids the aforementioned pitfalls, is preferable (Mahn et al., 2018).

Due to the close integration of active electrical circuitry in close proximity to recording electrodes, there remains the possibility that line noise, higher frequency electrical noise, or large voltage swings indicative of a floating ground may occur when the probe is connected to the OSC1lite. If Intan headstages are used, ensuring a solid ground/reference connection and separating the ground and reference by removing the 0 Ohm resistor on this headstage (where ground and reference are shorted by default) can greatly reduce or eliminate these noise issues. Additionally, while it is impossible to eliminate all stimulation artifact, selecting a waveform with a 1-2ms rise time can greatly reduce the offset observed during light onset/offset. For this reason, extra care must be taken when performing spike-sorting. Possible strategies for this include recording (via TTL events to the recording system) the onset and offset of each light pulse and removing these times from the recording file or, if the spike sorting software being used allows it, ignoring them during automatic clustering.

μLED probes are designed for focal neural perturbations and are well suited for interrogating neural microcircuits. However, these probes are not designed for targeting large brain volumes. While μLED probes can be used to verify the efficacy of one’s opsin in activating/inhibiting spiking in a given region, large-scale optogenetic perturbations are best performed with other light delivery systems.

Overall, μLED probes provide a powerful tool for interrogating neuronal circuits via simultaneous optogenetic perturbations and electrophysiology recordings from the same neurons.

## ACKNOWLEDGMENTS: (<u>Instructions</u>)

This work was supported by the following funding sources: NINDS 1F32NS117732-01, NSF 1545858, and NSF 1707316.

## DISCLOSURES: (<u>Instructions</u>)

E.Y. is a co-founder of NeuroLight Technologies, a for-profit manufacturer of neurotechnology. The remaining authors declare that they have no conflict of interest.

